# Modeling impacts of climate change on the potential habitat of an endangered Brazilian endemic coral: discussion about deep sea refugia

**DOI:** 10.1101/517359

**Authors:** Umberto Diego Rodrigues de Oliveira, Gislaine Vanessa de Lima, Paula Braga Gomes, Ralf Tarciso Silva Cordeiro, Carlos Daniel Pérez

**Affiliations:** Programa de Pós-Graduação em Ecologia, Universidade Federal Rural de Pernambuco, Recife, PE, Brazil.; Programa de Pós-Graduação em Biologia Animal, Universidade Federal de Pernambuco, Recife, PE, Brazil.; Departamento de Biologia, Universidade Federal Rural de Pernambuco, Recife, PE, Brazil.; Centro Acadêmico de Vitória, Universidade Federal de Pernambuco, Vitória de Santo Antão, PE, Brazil.

**Author notes:** These authors contributed equally to this work.

## Abstract

Climate and environmental changes are determinant for coral distribution and their very existence. Effects of such changes on distribution can be predicted through ecological niche models, anticipating suitable habitats for subsistence of species. *Mussismilia harttii* is one of the most widespread Brazilian endemic reef building corals, and in increasing risk of extinction. The ecological niche models were used through the maximal entropy approach to determine the potential present and future habitats for *M. harttii*, estimating suitable habitat losses and gains at the end of the 21st century. For this purpose, records published in the last 20 years and current and future environmental variables were correlated. The models were evaluated through the Area Under the Operational Curve of the Receiver, using the AUC values and additionally AUCratio, a new approach using independent occurrence data. Both approaches showed that the models performed satisfactorily in predicting areas of potential habitat for the species. The results showed that the area to the south of the São Francisco River is the most suitable for the current habitat of the species, and that nitrate was the most influential variable for the models. Simultaneously, the salinity and temperature exerted greater influence for the models in future scenarios, in which current northernmost and southernmost limits of the potential habitats shifted towards deeper regions, so these deeper sites may serve as a refugia for the species in global warming scenarios. Coral communities at such depths would be less susceptible to the impacts of climate change on temperature and salinity. However, deep sea is not free from human impacts and measures to protect deeper ecosystems should be prioritized in environmental policy for Brazilian marine conservation.

## Introduction

Coral reefs are one of the most valuable ecosystems on earth [1] providing a number of ecological services [2], such as shelter for associated fishes [3] and crustaceans [4, 5, 6, 7], also serving as substrate for coraline algae [8, 9]. Stable water conditions are determinant for the maintenance of living corals on reefs [10]. However, effects of climate changes put at least 50 % of shallow-water species in critical risk of extinction in the next 20 years [11, 12].

In the Southwestern Atlantic, coastal reef communities occur along of 3000 km of the Brazilian coastline [13], showing high endemism of reef-building species [14]. Four of those species belong to the genus *Mussismilia*, commonly known as brain-corals [15, 16, 17]. Although molecular assessments on *Mussismilia* are still rare [18], the distinctiveness between species is well established, allowing rapid identification on field [15]. The genus has at least two species in risk of extinction: *M. braziliensis* and *M. harttii* [19]. The first is restricted to shallow reefs of Bahia State and Abrolhos reefs, whereas the latter is found from the coast of Ceará to Espírito Santo States (from −3.822 to −40.583 latitud), commonly at depths of 2 - 6 m, and isolated records of up to 80 m [20].

*Mussismilia harttii* shows the lowest coverage percentages among its congeners [21], currently with populations in severe decline [19]. However, its conservation status at the IUCN (International Union for Conservation of Nature) database is still regarded as “Data Deficient” (DD). In contrast, the “Red Book of the Brazilian Fauna Threatened with Extinction” (2014, 2018), classifies the species as EN (Endangered) [19].

The distribution of marine organisms, such as corals, is determined by interactions of physical, chemical and biological factors [22]. Based on that, Ecological Niche Models (ENMs) approaches can provide information on the potential distribution of species within specific study areas [23]. ENMs associates environmental or spatial data to a set of distributional informations, such as distribution records [24], to outline the environmental conditions in which a given species may occur [25] indicating the most suitable areas for its occurrence [26, 27]. These models have been broadly applied to: prevent marine bioinvasions [28], conservation management planning [29], and especially to studies on climate changes [30, 31], predicting possible shifts on geographical distributions of key species [32].

The ENM also can be used to calculate the relative adequacy of a given habitat occupied by a species and to estimate changes in such suitability over time [33]. In the present study, we applied ENMs to generate maps of Current Potential Habitat (CPH) and Future Potential Habitat (FPH) for *M. harttii* at the end of the 21st century. These maps indicate potentially suitable areas and estimate habitat gains and losses in the different climatic scenarios projected. The projections will serve as tools for management plans and reef conservation in the southwest Atlantic reefs.

## Materials and methods

### Study area

The studied area comprises the Brazilian Exclusive Economic Zone - EEZ, in which *M. harttii* potentially occurs, from the intertidal zone to 100 m deep [34], based on all current records of its distribution [35, 36]. The study area also includes the priority areas for conservation according the Brazilian Ministry of the Environment (Portaria N° 19, of March 9, 2016 - ICMBio) (Fig 1).

**Fig 1.**
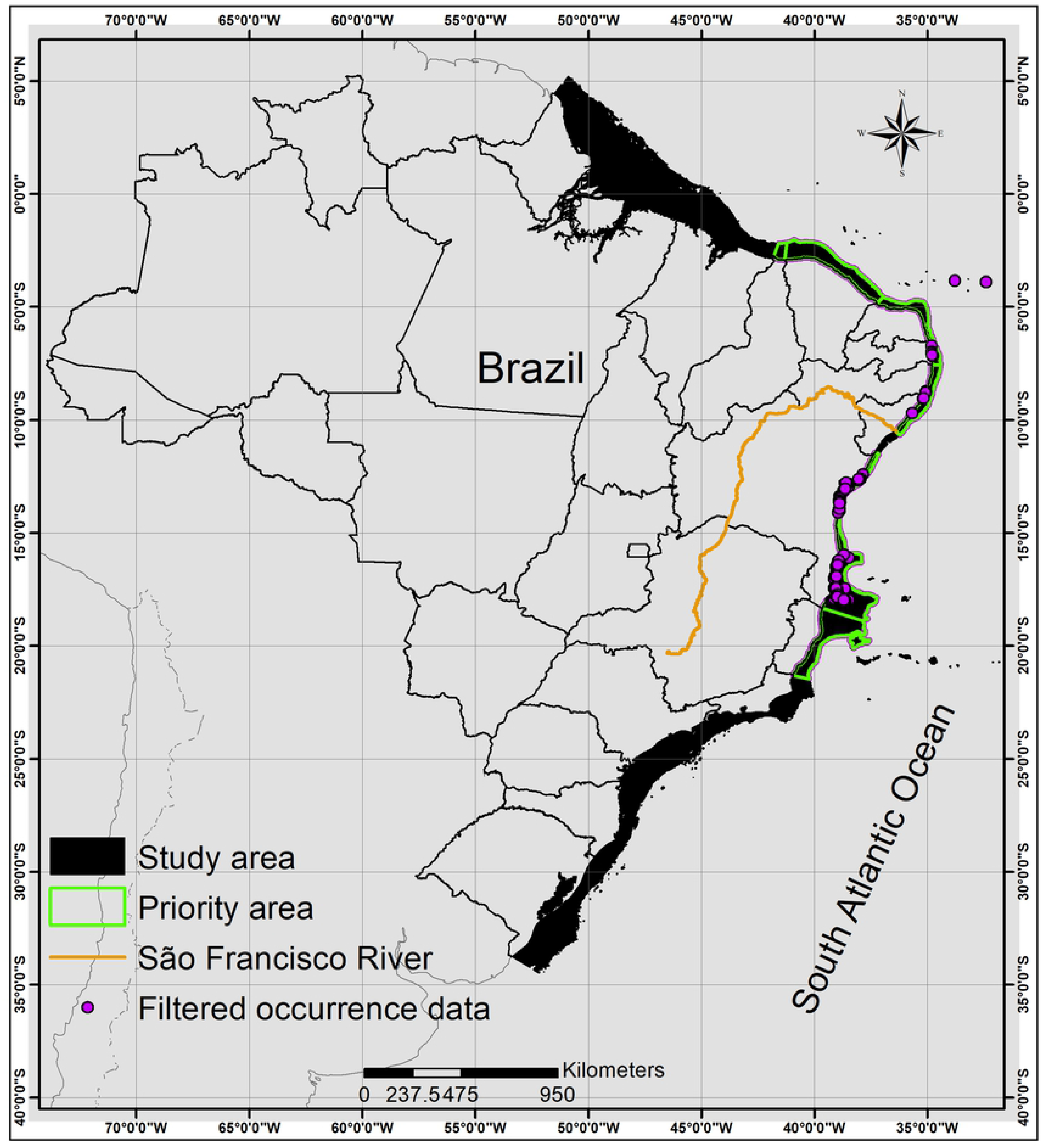
Map of the study area and occurrence records. Study area (Brazilian EEZ up 100 m), priority area for conservation of *M. harttii* and filtered occurrence data (one point in each pixel of 1km^2^).

### Occurrence records

An extensive search was made in specialized literature through academic indexing portals using the terms “Mussismilia”, “harttii”, “Brazil”, and “Brasil”, with publications containing precise geographic information (latitude, longitude and/or maps). Records of occurrence without georeferenced data were not used in the analyzes. These publication records were used to discuss the suitable area provided by the models. The search focused on records of *M. harttii* in the last two decades, avoiding the temporal decay in the quality of presence data due to the inherent dynamism of natural systems [37]. Sometimes, these data may be influenced by drastic phenomena, such as the local extinction of species [38] or changes in its distribution and abundance patterns [39]. Sampling bias on occurrence data are also common in areas of greater accessibility (more studied) because of regional interests [40]. This may reduce the model’s ability to predict the spatial data independence [41]. Alternatively, simple rarefaction method was used to reduce the autocorrelation of such points of occurrence, using SDMtoolbox v.2.2 [42], in which they were filtered (Fig 1), reducing data to only one point per pixel of 1km^2^, which was selected for modeling.

Species data collected *in situ*, from reefs located in the States of Paraíba, Pernambuco, Alagoas, Bahia, as well as independent species presence data, were not used during the modeling process, but *a posteriori* to evaluate the model [43].

### Selection of environmental layers

The environmental characterization variables provided by Bio-oracle available online (www.bio-oracle.org) were used. This global database provides current *in situ* and satellite-based oceanic information [22]. Bio-oracle also provides future variables based on the projections made by the International Panel on Climate Change (IPCC) for 2100 [44], in scenarios with different levels of greenhouse gas concentration (Representative Concentration Pathways - RCPs) [45]. These variables were cut to the extent of our study area [46] and re-sampled for the 30 arc seconds resolution (~ 1 km), as it is more indicated in local scale studies [47] and also due to low dispersal efficiency of the species.

The number of variables used may depend on the number of occurrence records [48], and when there are few records, such as endemic or threatened species, a small number of variables may be sufficient [49]. In order to generate the CPH map, 12 uncorrelated variables (Table 1) were selected through Pearson correlation matrix [50] with r < 0.8 (S1 Appendix), all ecologically or physiologically relevant [40].

**Table 1.** Details about the 12 Bio-Oracle variables used in the modeling process.

| Remotely sensed data | Variable | Sensor | Temporal range | Original spatial resolution | Unit |
| --- | --- | --- | --- | --- | --- |
|  | Calcite.mean | Aqua-MODIS § | 2002-2009 | 5 arc min (9.2 km) | mol / m<sup>3</sup> |
|  | Diffuse attenuation.max | Aqua-MODIS § | 2002-2009 | 5 arc min (9.2 km) | m<sup>-1</sup> |
|  | Temperature.max | Aqua-MODIS § | 2002-2009 | 5 arc min (9.2 km) | ° C |
|  | Photosynthetically available radiation.max | SeaWiF§ | 1997-2009 | 5 arc min (9.2 km) | Einstein / m<sup>2</sup> / day |
|  | Chlorophyll.range | Aqua-MODIS § | 2002-2009 | 5 arc min (9.2 km) | mg / m<sup>3</sup> |
|  | Primary productivity.max | PISCES | 2002-2009 | 5 arc min (9.2 km) | g / m<sup>-3</sup> / day<sup>-1</sup> |
|  | Present.surface.phytoplankton.min | PISCES | 2002-2009 | 5 arc min (9.2 km) | mmol / m<sup>-3</sup> |
|  | Current velocity.min | ORAP | 2002-2009 | 5 arc min (9.2 km) | m / s<sup>-1</sup> |
| In situ measured oceanographic data | Variable | Database | Temporal range | Number of data points | Unit |
|  | Salinity.max | World Ocean database 2009 † | 1961-2009 | 532377 | PSS |
|  | Dissolves oxygen.min | World Ocean database 2009 † | 1898-2009 | 540582 | ml / l |
|  | Nitrate.max | World Ocean database 2009 † | 1928-2009 | 189530 | µmol / l |
|  | pH | World Ocean database 2009 † | 1910-2007 | 117833 | - |

The projections of the IPCC for the year of 2100, developed by different research groups [51, 52], provide likely ranges of global temperature in future scenarios for population, economic growth and carbon use. These projections, called Representative Concentration Pathways (RCPs) [44], were used to model the *M. harttii* FPH in three different scenarios: decrease in emissions (RCP 2.6), stabilization of emissions (RCP 4.5) and increase in emissions (RCP 8.5) [53, 54, 55].

### Modeling process approach

The maximum entropy approach MaxEnt v. 3.3.3 [56, 57, 58] was used to model the potential distribution of *M. harttii*. MaxEnt is one of the most widely used algorithms for ENMs [59], because it presents consistent predictive performance compared to other algorithms [60], especially when the number of occurrence points is low [43, 61].

Table 1. Information on data searching and acquisition, and data availability of the 12 Bio-Oracle variables used in the modeling process, highlighting the five variables used to generate the Current Potential Habitat (CPH) model for *Mussismilia harttii*.

In the first step, we used relevant non-correlated variables and the filtered points of occurrence (Fig 1). The algorithm was calibrated using standard parameters [62], 1 % fixed omission threshold [63], 75 % of the records of occurrence for training and 25 % for test [64] (S2 Appendix), bootstrap (100 replicates) and maximum background number (5000). The Jackknife function of MaxEnt [65] was used to identify the percentage of contribution for each variable. In the second step, the five variables with the highest percentage of contribution (Table 1), the same points of occurrence and the same calibration of the MaxEnt were used to generate the CPH map.

To design the *M. harttii* FPH in the three future scenarios, MaxEnt was calibrated with the same parameters of steps 1 and 2, and also with: mean temperature, salinity and current velocity for the year 2100.

### Suitability area

Based on threshold values, the continuous maps of CPH and FPH were transformed into binary maps of suitability or probability [66], in which pixels are classified as “adaptive / presence” and “non-adaptive / absence”[43].

### Evaluation of the models

The Area Under the Receiver Operating Curve (AUC-ROC) is the most common metric to evaluate the accuracy of models [67]. AUC values ≤ 0.5 indicate that the model failed to perform better than random expectations, whereas values close to 1 indicate a good performance of the model [68]. In practice, the AUC-ROC is calculated based on a series of trapezoids [69], with the curve essentially “connecting the points” representing the different thresholds of the prediction [70]. This approach is used when input data is partitioned, in this case into training and test data [71]. When biotic data are divided into presence and absence (background), the AUC measures the discriminatory ability of the model to correctly predict the origin of these data if randomly selected [43].

Although the use of AUC-ROC for model evaluation is not questioned herein [72], we additionally used the partial ROC (AUCratio), an independent cutoff threshold metric where significant values are above 1 [73]. The AUCratio is a ratio between the predicted model AUC and null expectation [70] that a model generated with random data does not have a better prediction than the models generated with the input data [74]. We calculated the ratio of AUCrandon (at level of 0.5) and the AUCatual (calibrating 5% of omission and 500 bootstrap interactions) using the predicted distribution model [68] and independent occurrence records (S3 Appendix), through the package “ntbox” v.0.2.5.3 for Rstudio [75], to ensure greater robustness in model analysis [76].

## Results

One hundred and fifteen occurrence points were used for the modeling, of which 87 for training and 28 for testing and for external validation 24 points of occurrence were used. The variables with highest percentages of contribution and used to model CPH were, respectively: maximum nitrate (44.9 %); mean calcite (25.9 %); maximum salinity (21.3 %); maximum diffuse attenuation (5.8 %); and maximum temperature (2%).

The maximum training sensitivity plus specificity logistic threshold used to generate the binary maps maximized the sensitivity and specificity of the model [77]. This threshold is best suited for studies on rare or endangered species [74], as it reduces the over-prediction rate and selects only areas with high environmental suitability [43]. The thresholds of CPH (0.1391) and FPH (RCP 2.6- 0.1872, RCP 4.5- 0.1606 and RCP 8.5- 0.1702) show that a random prediction in a fraction of the same area does not have a better prediction than the points used in the test step [74].

The CPH of *M. harttii* represents a suitable area corresponding to 0.0418 % of the study area (Fig 2; Table 2). The sites north of the São Francisco River shows a smaller suitability (25,7 %) (Figs 2 a and b; Table 2), whereas the largest suitable areas are concentrated southwards of the São Francisco River (74,3 %) (Figs 2 c, d and e; Table 2). The AUC (S4 Appendix) and AUCratio (S5 Appendix) of the model were 0.979 and 1.934446, respectively.

**Fig 2.**
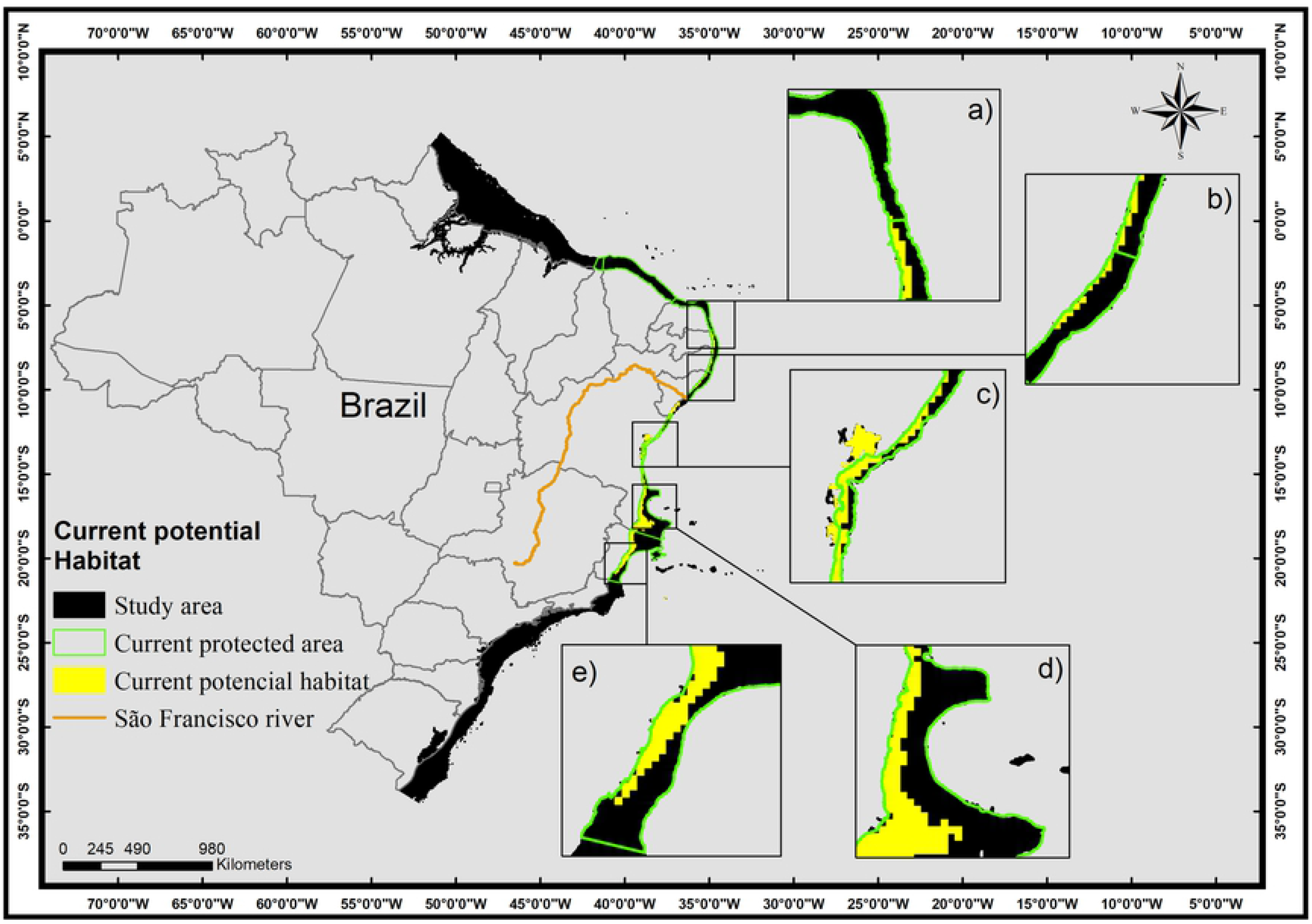
Map of Current Potential Habitat (CPH) of *Mussismilia harttii*. Highlighted figures (a, b, c, d, e) show concentrations of the highest number of suitable areas: a) Rio Grande do Norte and Paraíba States; b) Pernambuco and Alagoas States; c) north of Bahia State; d) south of Bahia State; and e) Espírito Santo State.

**Table 2.** Areas of suitable habitats

| Model | North area<br>(km<sup>2</sup>) | South area<br>(km<sup>2</sup>) | Total area<br>(km<sup>2</sup>) | New areas (km<sup>2</sup>) |  | Lost areas (km<sup>2</sup>) |  | Kept areas (km<sup>2</sup>) |  |
| --- | --- | --- | --- | --- | --- | --- | --- | --- | --- |
|  |  |  |  | North | South | North | South | North | South |
| CPH | 7951.5 | 22943.8 | 30895.3 |  |  |  |  |  |  |
| RCP 2.6 | 11720.8 | 38712.4 | 50433.2 | 8731.7 | 17403.5 | 962.4 | 5634.4 | 2989.1 | 21309.1 |
| RCP 4.5 | 13375.1 | 42987.6 | 56362.7 | 10132.1 | 20735.7 | 708.5 | 4691.5 | 3243 | 22252 |
| RCP 8.5 | 13581.3 | 43086.6 | 56667.9 | 10298.3 | 21545.7 | 668.4 | 5402.5 | 3283 | 21540.9 |

The three future distribution scenarios for *M. harttii* (RCP 2.6, RCP 4.5 and RCP 8.5) were characterized by an increase of suitable areas for the persistence of the species (67 - 88 %) (Table 2), but there was a significant reduction of suitable areas at the southern end of the distribution, at the Espirito Santo State (Figs 3, 4 and 5e). In all scenarios of FPH the salinity was the variable with the greatest contribution to the models (> 80 %), followed by temperature (~ 13 %) and current velocity (< 5 %).

**Fig 3.**
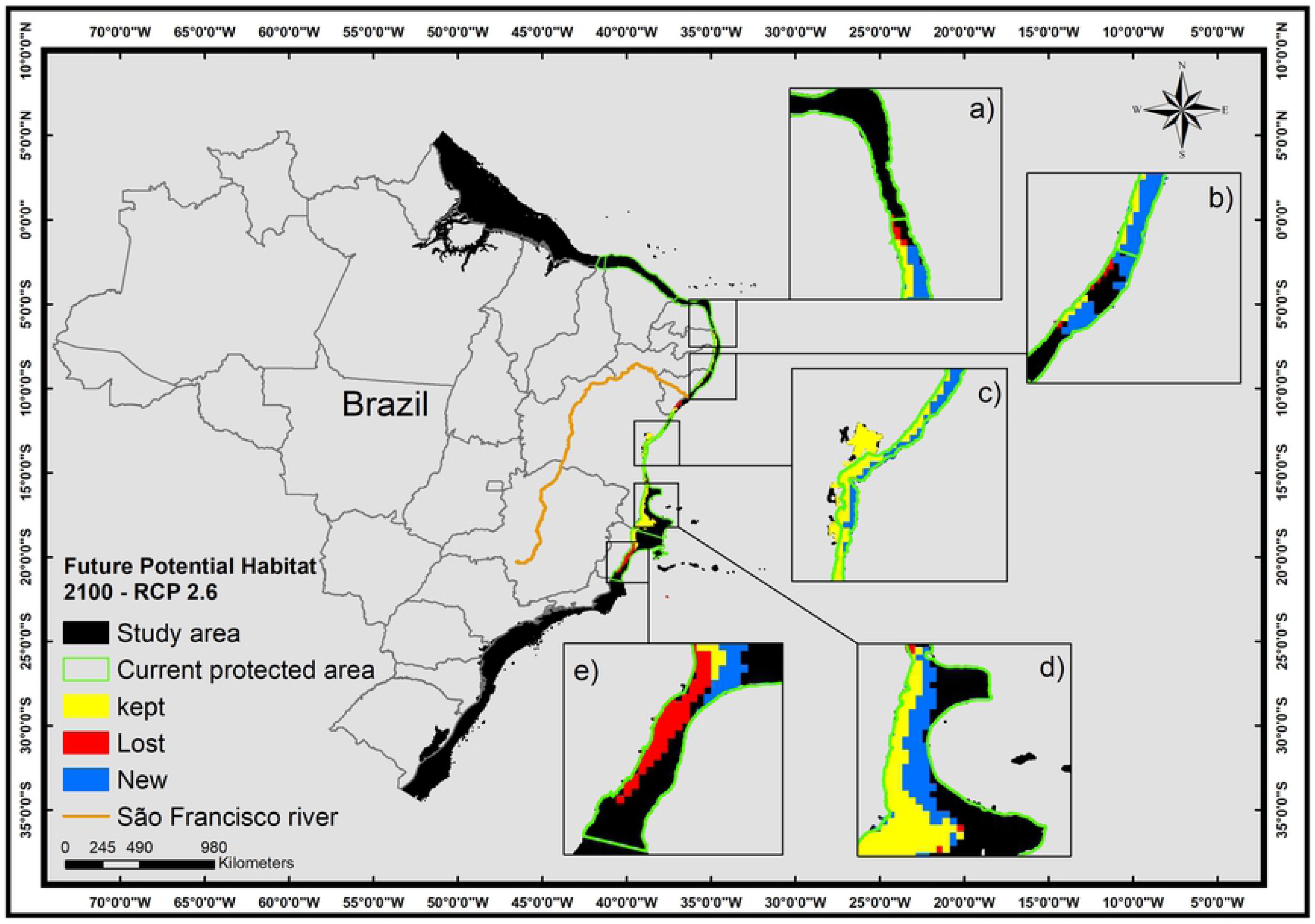
Map of Future Potential Habitat (FPH) of *Mussismilia harttii* in a scenario of reduction of greenhouse gas emissions (RCP 2.6) in the year 2100. FPH includes regions with kept, new, and lost suitability compared with the present (CPH). Highlighted figures (a, b, c, d, e) show concentrations of the highest number of suitable areas. a) Rio Grande do Norte and Paraíba States; b) Pernambuco and Alagoas States; c) north of Bahia State; d) south of Bahia State; and e) Espírito Santo State.

**Fig 4.**
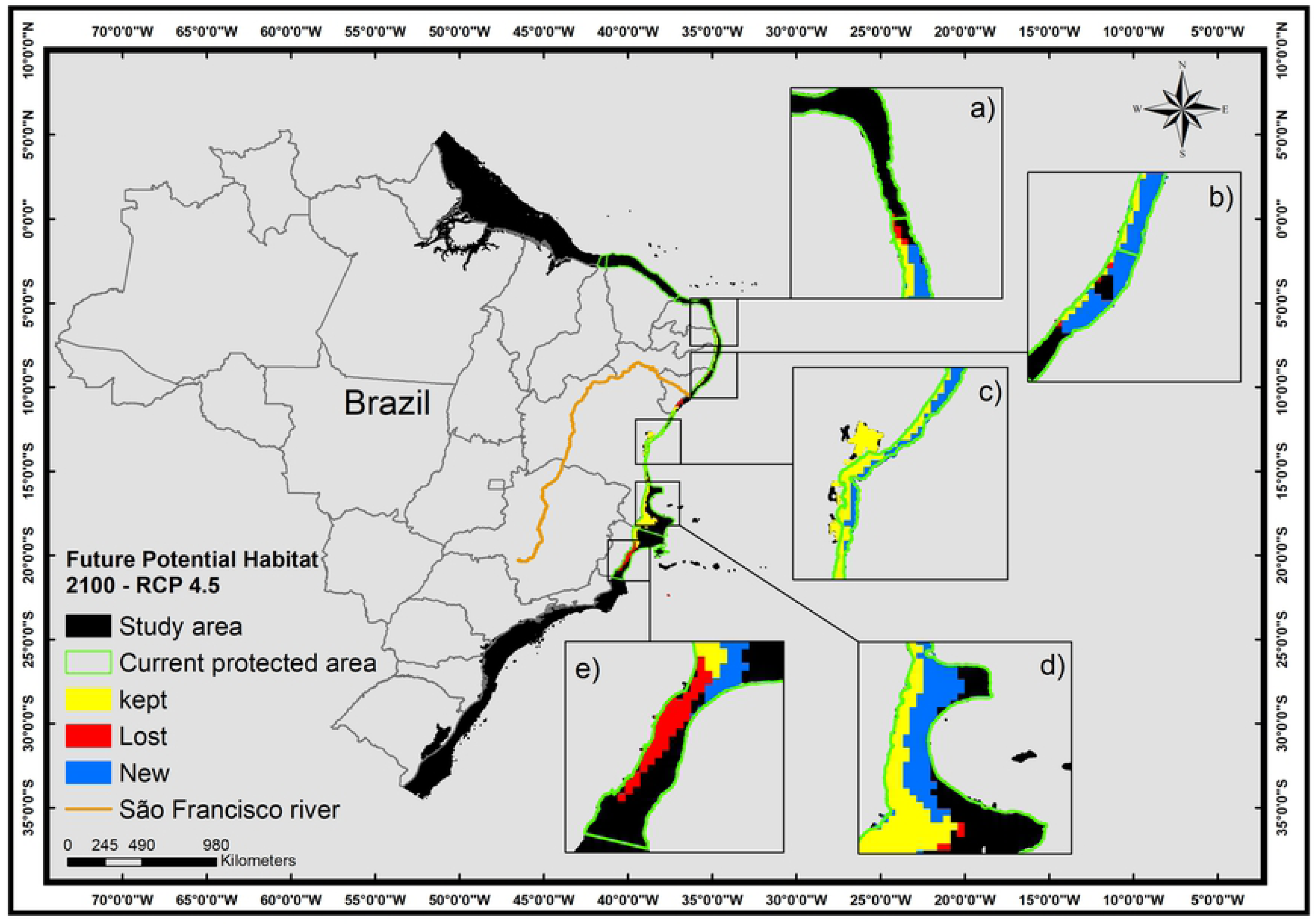
Map of Future Potential Habitat (FPH) of *Mussismilia harttii* in a scenario of reduction of greenhouse gas emissions (RCP 4.5) in the year 2100. FPH includes regions with kept, new, and lost suitability compared with the present (CPH). Highlighted figures (a, b, c, d, e) show concentrations of the highest number of suitable areas. a) Rio Grande do Norte and Paraíba States; b) Pernambuco and Alagoas States; c) north of Bahia State; d) south of Bahia State; and e) Espírito Santo State.

**Fig 5.**
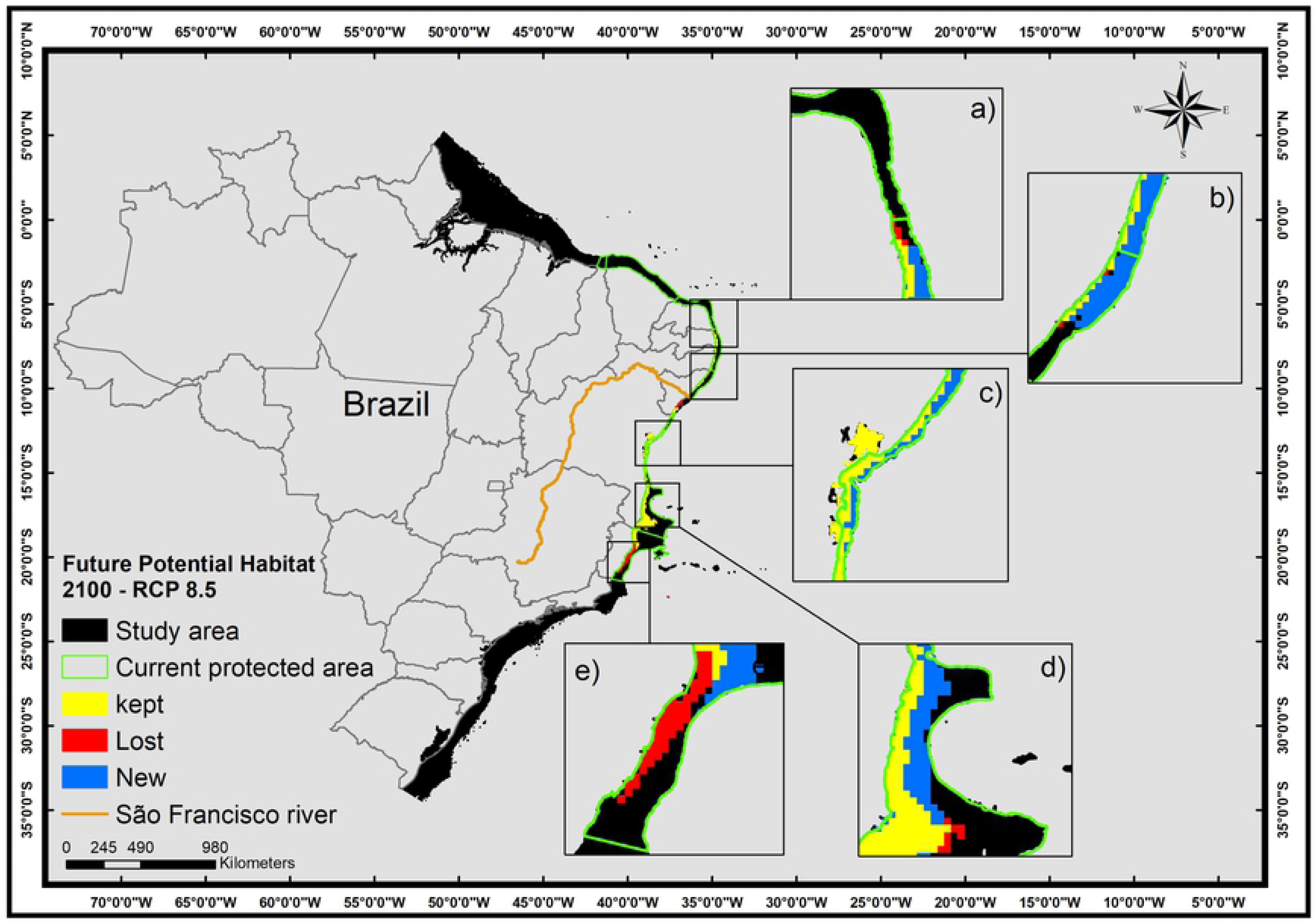
Map of Future Potential Habitat (FPH) of *Mussismilia harttii* in a scenario of reduction of greenhouse gas emissions (RCP 8.5) in the year 2100. FPH includes regions with kept, new, and lost suitability compared with the present (CPH). Highlighted figures (a, b, c, d, e) show the concentration of the highest number of suitable areas. a) Rio Grande do Norte and Paraíba States; b) Pernambuco and Alagoas States; c) north of Bahia State; d) south of Bahia State; and e) Espírito Santo State.

In a scenario of reduction of greenhouse gas emissions (RCP 2.6), the FPH of *M. harttii* represents a suitable area corresponding to 0.0805 % of the study area (Fig 3) (increasing 67 % of the CPH). The north of the São Francisco River shows a smaller area (23,2 %) (Figs 3a and b; Table 2), whereas the largest suitable areas are also concentrated to the south of the São Francisco River (76,8 %) (Figs 3c, d and e; Table 2). The AUC (S6 Appendix) and AUCratio (S7 Appendix) of the model were 0.975 and 1.914904, respectively.

In a scenario in which the emissions of greenhouse gases stabilize (RCP 4.5), the FPH of *M. harttii* represents a suitable area corresponding to 0.0881 % of the study area (Fig 4; Table 2) (increasing 87 % of the CPH). As with the previous scenarios, the sites northwards of the São Francisco River have a smaller suitable area (23.7 %) (Figs 4a and b; Table 2), while the largest areas of adequacy are concentrated southwards of the São Francisco River (76.3 %) (Figs 4c, d and e; Table 2). The AUC (S8 Appendix) and AUCratio (S9 Appendix) of the model were 0.973 and 1.912065, respectively.

In a scenario with increased greenhouse gas emissions (RCP 8.5), the FPH of *M. harttii* represents a suitable area corresponding to 0.0876 % of the study area (Fig 5; Table 2) (increasing 88 % of the CPH). The sites north of the São Francisco River again shows a smaller area (23.9 %) (Figs 5a and b; Table 2), whereas the largest suitable areas are concentrated southwards of the São Francisco River (76,1 %) (Figs 5c, d and e; Table 2). The AUC (S10 Appendix) and AUCratio (S11 Appendix) of the model were 0.973 and 1.911017, respectively.

Both current and future suitability areas for *M. harttii* are mostly within the Preservation Area for this species, with exception of Todos os Santos Bay, Bahia State (Figs 2, 3,4 and 5c). The three scenarios of future (year 2100) distribution of the species showed bathymetric expansion towards deeper areas, with a latitudinal restriction by the loss of suitable areas in the northernmost and southernmost limits of its distribution. (Figs 3, 4 and 5; Table 2).

In the current scenario (CPH), 60 % of the suitable areas are shallower than 20 m deep, 30 % between 20 - 50 m, and there are no suitable areas for the species north of the São Francisco River beyond 50 m. In the south, 0.87 % of the current suitable area corresponds to bathymetric ranges of 50 - 100 m (Table 3). In the three future scenarios (RCPs), 60 % of the new areas suitable for the species were concentrated between 20 m and 50 m, mostly to the south of the São Francisco River (Table 3).

**Table 3.** Areas (km^2^) of suitable habitats by depth ranges

| Model | Depth | 0 - 20 m | 20 - 50 m | 50 - 75 m | 75 - 100 m |  |  |  |  |
| --- | --- | --- | --- | --- | --- | --- | --- | --- | --- |
| CPH | North | 3155.1 | 458.8 | 0 | 0 |  |  |  |  |
|  | South | 14635.8 | 8567.9 | 189.4 | 16.1 |  |  |  |  |
|  | Lost areas (km<sup>2</sup>) |  |  |  |  | New areas (km<sup>2</sup>) |  |  |  |
|  | Depth | 0 - 20 m | 20 - 50 m | 50-75 m | 75 - 100 m | 0 - 20 m | 20 - 50 m | 50 - 75 m | 75 - 100 m |
| RCP 2.6 | North | 641.6 | 174.2 | 0 | 0 | 1519.8 | 7935.2 | 1789.7 | 1123.9 |
|  | South | 2924.7 | 3881.1 | 290.6 | 75.2 | 5155 | 15078.9 | 3854.8 | 1992.5 |
| RCP 4.5 | North | 687.7 | 150 | 0 | 0 | 1987.7 | 9168.7 | 2010.2 | 1271.7 |
|  | South | 1923 | 3679.7 | 374.1 | 75.2 | 5786 | 18209.2 | 4855.1 | 2550.3 |
| RCP 8.5 | North | 772.6 | 108.4 | 0 | 0 | 2087.3 | 9369.7 | 1983.1 | 1247.5 |
|  | South | 2428.2 | 4132.2 | 282.6 | 22.6 | 5473.7 | 18523.6 | 5495.7 | 2516 |

In summary, in the future scenarios there was a latitudinal restriction of appropriate areas for *M. harttii* (- 6.751 ° to - 19.894 ° latitud), but it increased (67 - 88 %) towards deeper waters.

## Discussion

### Visualization and Interpretation of Ecological Niche Models

Predicting and mapping potential suitable habitats for threatened and endangered species is critical for monitoring and restoring their natural populations [78]. In this sense, the modeling approach is an effective tool, which can predict the direction of contractions and expansions of species distribution [79], producing probability maps for presence or relative suitability of a species [80].

Table 3. Values of current potential habitat areas (CPH) and future potential habitat areas for *Mussismilia harttii* in three different projected climatic scenarios for the year 2100 (RCP 2.6, RCP 4.5 and RCP 8.5), north and south of the São Francisco River, arranged in four depth ranges.

Besides elevated CPH validation indexes (AUC and AUCratio were 0.979 and 1.934446, respectively), literature data (not geo-referenced and therefore not used in the model) also record *M. harttii* in areas indicated by the model as suitable for the species, such as the southern portion of the Abrolhos bank at Espírito Santo State [81] and at coastal reefs of Rio Grande do Norte State [82]. A model that fails to omit known points of presence is more flawed than those that predict unknown inhabited areas [83]. These unknown areas do not represent a model prediction error, but provide a precise representation of the spatial extent of habitable conditions for the species [70].

The area of potential species habitat is generally larger than the real distribution [56] and projections beyond the time interval of a training dataset (distribution in future dates) require cautious interpretations to avoid possible misinterpretations [84]. Such caution is because AUC values tend to increase when the selected background area is larger than the observed current habitat of a species [85]. Although the AUC values (close to 1) showed that the models performed very well with the results [77, 78] (better than any model generated with a set of random predictors [71], it was necessary to use a different approach metric for evaluate the models. In the AUC metric, the weight of commission errors is much lower than that of omission errors, which makes it an inappropriate performance measurement [86].

The AUCratio also showed a good performance of the model, with values above 1 [70] and close to 2. These results allowed us to evaluate the statistical significance of the AUC itself [86]. In this way, it is more appropriate to evaluate the model performances [72]. The thresholds used to generate the binary maps are best suited for applications in Ecological Niche Templates [87] by better predicting independent occurrence data [46].

### Environmental variables and *M. harttii’s* habitat

Even though the effects of each environmental variable over the population dynamics are unknown [88], the variables chosen to model the habitat suitability for *M. harttii* are in accordance with default conditions in previous studies on anthozoans [62]. Nitrates are the most common form of dissolved inorganic nitrogen in coastal waters, being the main contributor for the CPH [89]. Long exposure to high nitrate levels may lead to bleaching in some corals, due to zooxanthellae loss, on the other hand, in high temperatures, the nitrate enhancement may sustain the remaining zooxanthellae for a short period until their reestablishment, as a compensatory mechanism [90]. Calcite is one of the most common forms of calcium carbonate [91], and it was the second most important variable for the CPH. Studies indicate that calcification ratios in tropical reef-building corals will be reduced in 20 - 60 %, when CO2 concentrations reach twice the pre-industrial concentration levels (around 560 ppm) [92].

Future habitat scenarios for *M. harttii* were mostly influenced by salinity and temperature. However, shifting of suitable habitats to deeper areas can be related with several factors. Future climate projections show not only a temperature increase of the ocean. Temperature increase will affect regimes of winds, ocean circulation and, consequently, precipitation and continental runoff, which directly influences the salinity in coastal waters [93, 94]. As result, higher turbidity and lower salinity are expected in such areas. Despite Brazilian corals as a whole are considered resistant to the input of terrigenous sediments [88], *M. harttii* has preference for clear waters, in which it is more abundant than in turbid zones [95]. Typically, corals dwell habitats under salinities between 32 and 40 [96]. Fall in salinity, even in short term, may lead to reduction of fertility [97], increase of susceptibility to bacterial infections [98], being potentially lethal to corals and their endosymbionts [99, 100].

Temperature, salinity and light have major effects on where reef-building corals grow [100]. Despite the temperature showed the lowest contribution for the CPH, it is undoubtedly determinant for the future persistence of coral species, as 50% of these corals are threatened by climate changes [11, 12]. Our results also show the importance of the temperature in the FPH for *M. harttii*. This species suffer thermal stress in temperatures higher than 31.0°C, leading to long-term damage or death [101]. In fact, a recent study reported massive coral bleaching events in temperatures above 27° in Abrolhos reefs [102], which concentrate most records of *M. harttii* in the present study (fig 1).

Another important factor is the competition with algae (macroalgae and filamentous algae). A recent study on Brazilian benthic communities showed that such organisms dominate reefs down to 15 m deep [103]. Algae are favored by anthropic impacts, such as reduction of herbivorous/grazer fishes by overfishing, and increase of nutrients from land [104, 105]. Thus, in future scenarios, algae will likely continue to be favoured, and its competition with corals tend to reduce coverage of the later in shallow waters. In contrast, deeper areas would be less susceptible to the influence of runoff, temperature and salinity changes. Despite the lack of earlier baselines for Brazilian benthic communities, it is possible to affirm that the current scenario is result of a sum of anthropic impacts, as studies back in the 1960’s describe distinctive zonation and coverage in these communities [106].

### Current distribution of *M. harttii*

Most of the current suitable distribution area for *M. harttii* (CPH) is southwards of the Rio São Francisco river, where most published records are concentrated. Despite records in the coast of the Espírito Santo State (~ 19 ° S) were absent in our analyses, that area is known as the southernmost distribution limit for the species [95], with the highest percentage of CPH. That region coincides with a center of diversity within the Brazilian Province (20 ° S to 23 ° S), as indicated for benthic organisms, such as algae, invertebrates and fishes [103; 107; 108; 109]. That center is favored by the confluence of currents in the Brazilian coast, creating a transition zone between tropical and subtropical diversity [103]. Despite a limited number of records of *M. harttii* and a smaller percentage of CPH to the north of the São Francisco river, the species is the main reef-builder northwards the São Francisco river [14].

Most records of *Mussismilia harttii* are from shallow reefs, between 2 and 6 meters [106] and consequently close to the coast. However, scattered records show this species occupying deeper reefs (up to 25 m) [81] and even at mesophotic depths [20]. Similarly, most of the CPH is concentrated in shallow waters (0 - 20 m), but with deeper suitable habitats commonly occurring, especially in the southern portion of species distribution.

### *Mussismilia harttii*’s response to climate change by the end of the 21st century

Future distribution models (RCP 2.6, RCP 4.5 and RCP 8.5) of *M. harttii* showed expansion of suitable areas, in relation to the current habitat, towards deeper sites where there are few records of this species. Concomitantly, there was a reduction of suitable shallow water areas, especially at the southernmost distribution limit, which suffered the greatest losses (Fig 3e, 4e and 5e). A similar effect is expected in the same are (mainly in the Espirito Santo State), as previous ENMs studies also showed losses for the zoanthid *Palythoa caribaeorum* [62].

A recent study on *M. harttii* [82] estimates a decline of its populations in their current geographic range in shallow waters. Our results also indicate an future scenario (RCP 8.5) with a loss of 25 % of the current suitable area (7,746.6 km^2^ lost) in shallow waters (0 - 20 m), concentrated mainly in the southernmost distribution of the species (Espírito Santo State) (Table 2). Conversely, the results show a 55 % increase at deeper areas, 20 - 50 m (Table 3). Thus, in a future scenario, the species would lose suitable habitats in coastal shallow sectors, followed by a gain of deeper habitats, which could serve as refugia in face of climate changes.

### Deep sea refugia strategy

The “deep reef refugia hypothesis” (DRRH) considers that coastal anthropic impacts and thermal stress effects are progressively reduced with depth [110, 111, 112]. Therefore, mesophotic coral ecosystems, between 30 and 150 m, have been treated as important refugia for shallow reefs diversity [113; 114], temporarily supporting coral populations from shallow-reefs under stress conditions [115]. Such areas would provide shelter in which these populations might persist and from which would subsequently expand [116], recovering previously damaged areas [103, 117].

The reduction of shallow suitable areas and increase of deeper habitats suggest the potential of *M. harttii* for using mesophotic reefs as refugia, ensuring its subsistence. However, the DRRH is more adequate for species with wide depth distribution ranges [103] and presupposes larvae exchange between deep and shallow populations [118], which have been demonstrated to be local and species-specific [119]. Despite *M. harttii* is particularly representative in shallow waters (2-6 m), scattered records show this species occupying deeper reefs (up to 80 m) [20, 81, 120] (S12 Appendix). which reinforces the potential of the species to occupy deep mesophotic areas.

Even showing wide depth ranges, connectivity between coral populations is not always continuous along bathymetric gradients [121]. Consequently, it is still unknown if deeper populations of *M. harttii* would serve as genetic stocks for shallow waters, as most of its deep records are sparse and rare [111, 118]. In any case, the expansion of deeper suitable areas may result in the expansion of deeper populations of *M. harttii*, regardless of the maintenance of coastal populations. In case of connectivity, such refugia would contribute for the recolonization of the coastal zone affected.

Studies using of global climate models mostly suggest that few shallow coral species will persist under a sea surface temperature increase of 2 ° C in the next one hundred years [122]. Nevertheless, given the current slowness in mitigation measures, it is expected an increase of 3,1 ° C in the same period (RCP 8.5) [123]. In such scenarios, identify and protect deep sea refugia must become a priority for species conservation [114].

### Threats and perspectives for conservation

The main global threats to coral species are related with greenhouse gas emissions (RCP), especially CO_2_ [104]. Effects of such impacts have lead to decline of biodiversity in reefs of Brazil and of the world, through increase of sea temperature and ocean acidification [11]. Local impacts boost these effects through higher sedimentation, multiple biological invasions, bleaching, coral diseases and, consequently, loose of diversity on reef environments [11, 124, 125, 126]. Such impacts are frequently related to disorganised urban growth, pollution, messy tourism practices and overfishing [127, 128, 129]. In the literature *Mussismilia harttii* used to be described as forming extensive bands on coastal reefs, showing colonies usually up to 1 m in diameter [106]. Currently, this is a rare scenario for most of these reefs, which often have a low coral coverage, not corresponding the descriptions of the 1970’s.

Environmental changes have triggered reorganisations in reef ecological relationships, zonation and dominance, in processes also called *phase-shifts* [130]. In most reefs, for example, scleractinian dominance have been replaced mainly by macroalgae [105], octocorals [131], sponges [132] and/or zoanthids [133, 134, 135], the latter is the case of the Brazilian reefs [136]. In these reefs, *M. harttii* is also threatened by the dominance of invasive species, such as *Tubastraea* spp. [137], which even more compromises its resilience of shallow reefs.

The accelerated loss of biodiversity and habitats is one of the worst crisis of the present time, as evidenced by the ever increasing species red lists. All current and future scenarios showed herein alert for the relevance of the endemism and the role of *M. harttii* as a reef builder in Brazilian reefs. Currently, the species is classified as “in risk of extinction” [19], and the perspective of reduction of suitable shallow areas highlight the urgency of priority conservation measures. Future environmental politics, therefore, must focus not only in the recover of coastal populations, but also on the conservation of mesophotic coral ecosystems (MCE’s). Despite being less affected by climate changes, MCE’s are impacted by human activities, such as fisheries, mining and drilling [119, 138] and measures to protect deeper ecosystems should be prioritized in environmental policies for marine conservation, especially in Brazil.

## Supporting information

**S1 Appendix. Pearson correlation matrix of environmental variables**. Pearson correlation matrix of 39 environmental variables, in which the 12 variables with no correlation greater than 0.8 were chosen.

**S2 Appendix. Occurrence records used to generate maps of Current Potential Habitat and Future Potential Habitat.** Georeference (latitude and longitude), source and author of occurrence records used to generate the CPH and FCP models.

**S3 Appendix. Occurrence records used to evaluate models through AUCratio**. Georeference (latitude and longitude) and author of occurrence records used to evaluate the models through AUCratio.

**S4 Appendix. Maxent output of Current Potential Habitat**. Maxent output with values of threshold, AUC, percentage of the predicted area and number of occurrences used to generate the Current Potential Habitat model.

**S5 Appendix. Output of the ntbox used to evaluate the AUCratio of CPH**. Values of AUCratio for a AUC_randon_ (at level of 0.5) and the AUC_atual_ (calibrating 5% of omission and 500 bootstrap interactions).

**S6 Appendix. Maxent output of FPH (RCP 2.6)**. Maxent output with values of threshold, AUC, percentage of the predicted area and number of occurrences used to generate the FPH model (RCP 2.6).

**S7 Appendix. Output of the ntbox used to evaluate the AUCratio of FPH (RCP 2.6)**. Values of AUCratio for a AUCrandon (at level of 0.5) and the AUCatual (calibrating 5% of omission and 500 bootstrap interactions).

**S8 Appendix. Maxent output of FPH (RCP 4.5)**. Maxent output with values of threshold, AUC, percentage of the predicted area and number of occurrences used to generate the FPH model (RCP 4.5).

**S9 Appendix. Output of the ntbox used to evaluate the AUCratio of FPH (RCP 4.5)**. Values of AUC_ratio_ for a AUC_randon_ (at level of 0.5) and the AUC_atual_ (calibrating 5% of omission and 500 bootstrap interactions).

**S10 Appendix. Maxent output of FPH (RCP 8.5)**. Maxent output with values of threshold, AUC, percentage of the predicted area and number of occurrences used to generate the FPH model (RCP 8.5).

**S11 Appendix. Output of the ntbox used to evaluate the AUCratio of FPH (RCP 8.5)**. Values of AUCratio for a AUC_randon_ (at level of 0.5) and the AUC_atual_ (calibrating 5% of omission and 500 bootstrap interactions).

**S12 Appendix. Unpublished work**. Cordeiro, RTS; Amaral, FMD. Ocorrência de cnidários construtores de recifes em ambientes de profundidade no Nordeste do Brasil. In: Abstracts of XIV Congreso Latinoamericano de Ciencias del Mar, 2011, Balneário Camboriú - SC, Brazil.

## Acknowledgments

We thank who collaborate with advising and collection of data used herein: Maude Gauthier (Sherbrooke University), Catherine George (Sherbrooke University), Manuela Menezes, David Montenegro, Rafael Brandão, Erika Santana e David Oliveira. Gratitude to all boatmen, *jangadeiros* and fishermen of Northeast from Brazil, which make this work possible. This study is a product of the Anthozoan Research Group - GPA (UFPE-UFRPE).

